# Glutamatergic dysfunction leads to a hyper-dopaminergic phenotype through deficits in short-term habituation: a mechanism for aberrant salience

**DOI:** 10.1101/2021.07.23.453593

**Authors:** Marios C Panayi, Thomas Boerner, Thomas Jahans-Price, Anna Huber, Rolf Sprengel, Gary Gilmour, David J Sanderson, Paul J Harrison, Mark E Walton, David M Bannerman

## Abstract

Psychosis in disorders like schizophrenia is commonly associated with aberrant salience and elevated striatal dopamine. However, the underlying cause(s) of this hyper-dopaminergic state remain elusive. Various lines of evidence point to glutamatergic dysfunction and impairments in synaptic plasticity in the aetiology of schizophrenia, including deficits associated with the GluA1 AMPAR subunit. GluA1 knockout (*Gria1*^-/-^) mice provide a model of impaired synaptic plasticity in schizophrenia and exhibit a selective deficit in a form of short-term memory which underlies short-term habituation. As such, these mice are unable to reduce attention to recently presented stimuli. In this study we used fast-scan cyclic voltammetry to measure phasic dopamine responses in the nucleus accumbens of *Gria1*^-/-^ mice to determine whether this behavioral phenotype might be a key driver of a hyper-dopaminergic state. There was no effect of GluA1 deletion on electrically-evoked dopamine responses in anaesthetized mice, demonstrating normal endogenous release properties of dopamine neurons in *Gria1*^-/-^ mice. Furthermore, dopamine signals were initially similar in *Gria1*^-/-^ mice compared to controls in response to both sucrose rewards and neutral light stimuli. They were also equally sensitive to changes in the magnitude of delivered rewards. In contrast, however, these stimulus-evoked dopamine signals failed to habituate with repeated presentations in *Gria1*^-/-^ mice, resulting in a task-relevant, hyper-dopaminergic phenotype. Thus, here we show that GluA1 dysfunction, resulting in impaired short-term habituation, is a key driver of enhanced striatal dopamine responses, which may be an important contributor to aberrant salience and psychosis in psychiatric disorders like schizophrenia.

## INTRODUCTION

Psychosis is a key feature of several neuropsychiatric disorders, including schizophrenia. Current thinking posits that psychosis is a disorder of aberrant salience [1–3]. Aberrant salience describes when a stimulus grabs inappropriately high levels of attention and thus drives maladaptive behavior. Aberrant salience is likely mediated via elevated dopamine (DA) levels, which have been strongly implicated in schizophrenia [3, 4]. However, the underlying causes of this DA dysregulation are often unspecified [5]. Although this hyper-dopaminergic state and aberrant salience could stem from primary abnormalities within the DA system, it could also constitute a final common pathway [5], resulting as a secondary consequence of other brain disturbances like glutamatergic dysfunction and impairments in synaptic plasticity, which are also strongly linked to schizophrenia [6–8]. Nevertheless, direct evidence for this hypothesis has remained elusive.

Recent large scale GWAS meta-analyses of schizophrenia have revealed an over-representation of genes associated with glutamate synapses and synaptic plasticity. For example, a genome-wide significant association to schizophrenia has been established for the Gria1 locus which codes for the GluA1 subunit of the AMPA glutamate receptor [9, 10]. GluA1 plays a key role in AMPAR trafficking and thus can support the expression of long-term potentiation (LTP), an experimental model of synaptic plasticity. Post-mortem studies have also demonstrated that GluA1 protein and mRNA expression are decreased in patients with schizophrenia (e.g. [11–15], which are unlikely to be attributed to long term neuroleptic treatment e.g. [16–19]). Together, these studies point to an important role for GluA1 in the aetiology of the disorder. *Gria1*^-/-^ mice therefore represent an important model of impaired synaptic plasticity in schizophrenia.

*Gria1*^-/-^ mice lack a form of short-term memory which results in deficits in short-term habituation [20–24]. As such, *Gria1*^-/-^ mice are unable to reduce the attention paid to a recently presented stimulus [25]. For example, whereas wild-type animals will reduce orienting and exploration, and pay less attention to a specific stimulus they have just experienced, *Gria1*^-/-^ mice fail to habituate and continue to pay attention to the stimulus when experienced again soon after [23, 24, 26]. This abnormally persistent attention to stimuli has been proposed as a model of aberrant salience [27]. However, the relationship between GluA1, short-term habituation and striatal DA release is unknown. Here we address this by recording phasic DA signals in the nucleus accumbens of wild-type and *Gria1*^-/-^ mice using fast-scan cyclic voltammetry (FSCV). We show that electrically evoked DA in anaesthetized *Gria1*^-/-^ mice is comparable to controls. However, phasic DA signals in response to both sucrose rewards and neutral light stimuli fail to habituate in awake behaving Gria1^-/-^ mice, resulting in a behaviourally relevant, hyper-dopaminergic phenotype.

## METHODS

See **<u>Supplementary Information</u>** for detailed methods.

### Subjects

Genetic construction, breeding, and genotyping of Gria1^-/-^ mice was as previously described [28]. Briefly, homozygous knockout (*Gria1*^-/-^) and wild-type (WT) littermates were generated from heterozygous-heterozygous breeding pairs. All procedures were performed in accordance with the United Kingdom Animals (Scientific Procedures) Act of 1986 and were approved by the University of Oxford Ethical Review Board. Subjects for anaesthetised voltammetry recordings were 6 WT mice (Male n=3) and 6 *Gria1*^-/-^ mice (Male n=3). During freely moving in-vivo voltammetry, data were obtained from N = 25 working electrodes (WT n=16, *Gria1*^-/-^ n=9) targeting the nucleus accumbens (NAcc) in N=14 mice (Male WT n=9, Male *Gria1*^-/-^ n=5).

### Fast scan cyclic voltammetry in anaesthetised recordings

Carbon fibre electrodes were constructed as described previously [29–31]. Those used in anaesthetised recordings were pre-calibrated using a flow cell to establish a conversion between current signals (nA) and DA concentrations (nM). Mice were surgically implanted with a bipolar stimulating electrode targeting the ventral tegmental area (VTA) and a carbon-fibre voltammetry electrode targeting the NAcc, and maintained under urethane anaesthesia for the recording procedure.

To establish whether GluA1-receptor knockout alters electrically-evoked DA release, we tested the following stimulation parameters: (1) Baseline responses (6 × baseline stimulations with 5 mins interstimulus interval (ISI)); (2) Stimulation intensity which was varied at 50, 100, 150, 200, 250, 300 μA; 2 measurements per stimulation amplitude, in ascending intensity order with a 3 min ISI; and (3) Number of pulses (5, 10, 20, 30, 40; 2 measurements per stimulation amplitude, in ascending intensity order with a 3 min ISI). (4) Finally, a second baseline of 6 responses was taken to assess the stability of the evoked response.

### Fast Scan Cyclic Voltammetry in Freely Moving Animals

Mice were implanted with bilateral carbon fibre voltammetry electrodes targeting the NAcc. Following post-operative recovery, mice were food restricted and maintained >85% of their free feeding weight. Mice were first acclimatized to the operant chambers (Med Associates), to the liquid sucrose reward and to the voltammetric headstage. They were then trained to nose poke in the food magazine. 22 μL sucrose reward was delivered upon magazine entry on a variable interval (VI) 60s and 120s schedule for 3 days each. Each session lasted 30 mins.

During two recording sessions, in addition to the VI120s nose poking schedule for sucrose, the mice were given non-contingent presentations of one of two different neutral visual stimuli lasting 10s each (comparable to [24]). One stimulus consisted of two flashing (0.909Hz) LED lights positioned either side of the reward magazine (‘LED’) and the other was the continuously illuminated house light on top of the conditioning box (‘House’). Presentation of lights occurred independently of the reward delivery schedule enabling an independent analysis of reward and stimulus evoked DA. To assess stimulus-specific short-term habituation of DA signals, each session consisted of 8 pairs of stimulus presentations (a pair of stimulus presentations constitutes a trial), with the delay between 1st and 2nd stimulus presentations in each pairing being 30s and the inter-trial interval being 310s (based on [24]). The trials in a session consisted of a pseudorandom permutation of the 4 possible pair types: House->House, LED->LED (both *“same”* trials), House->LED or LED->House (both *“different”* trials). If the DA response habituated in a stimulus-specific manner, then it would be smaller to the presentation of the identical cue as the second target stimulus on same trials than to the alternative cue as the second target stimulus on different trials.

In separate sessions, the mice were presented with a variable reward task. Rewards were again delivered contingent on a nose poke into the food magazine on a VI120s schedule, but now the magnitude of the sucrose reward was pseudorandomly varied (11, 22, and 44 μL; small, medium, and large rewards respectively).

### Histology

At the end of experimental procedures, mice were deeply anesthetized with pentobarbital (200mg/kg), and microlesions were made at the working electrode locations via current stimulation. The animal was transcardially perfused with saline followed by 10% formalin solution. Brains were sectioned in 40 μm coronal slices using a cryostat and were stained with Cresyl violet allowing histological identification of the electrode location.

## RESULTS

### GluA1 deletion does not alter sensitivity to electrically evoked VTA-NAcc DA release in anaesthetized mice

We first wanted to assess whether there were intrinsic differences in the strength of the mesolimbic pathway, in the releasable pool of DA, and in the kinetics of DA transmission in NAcc in the *Gria1*^-/-^ mice. To do this, we examined DA release following electrical stimulation of the VTA in anaesthetized mice (Fig 1A,B; Supplementary Fig 1A). Following stimulation, there were no significant genotype differences in the release and reuptake of DA. Specifically, there were no significant genotype differences on peak DA release (t_10_ = .42, *p* = .68; Fig 1C), latency to peak (t_10_ = 1.55, *p* = .15; Fig 1D), or rate of decay (t_50_ from model fit of negative exponential decay for traces with model fit R^2^> 90%; t_10_ = .32, *p* = .76; Fig 1E). Given that release and reuptake parameters are closely related, we used a multivariate ANOVA to directly compare these three outcome measures (peak DA, latency to peak, and rate of decay) to ensure there were no significant genotype differences in some simultaneous combination of these parameters. There was no significant effect of Genotype (F_1,10_ = .17, *p* _=_.69), or interaction between Genotype and the three outcome measures (Hotelling’s Trace (2,9) = .44, *p* = .66). Furthermore, there was no evidence of any genotype differences in the peak electrically evoked DA levels across a number of pulses (Fig 1F), or a range of stimulation intensities (Fig 1G; all F’s < 1.57; p > 0.22; Supplementary Fig 1B,C and Supplementary Results). Surprisingly, there was a significant sex difference such that peak electrically evoked DA was higher in male than female mice. However, this sex difference did not interact with Genotype (Supplementary Fig 1C,D). Together, this demonstrates that GluA1 deletion does not lead to intrinsic differences in electrically-evoked release or the balance of DA release/reuptake in this mesolimbic DA pathway.

**Figure 1.**
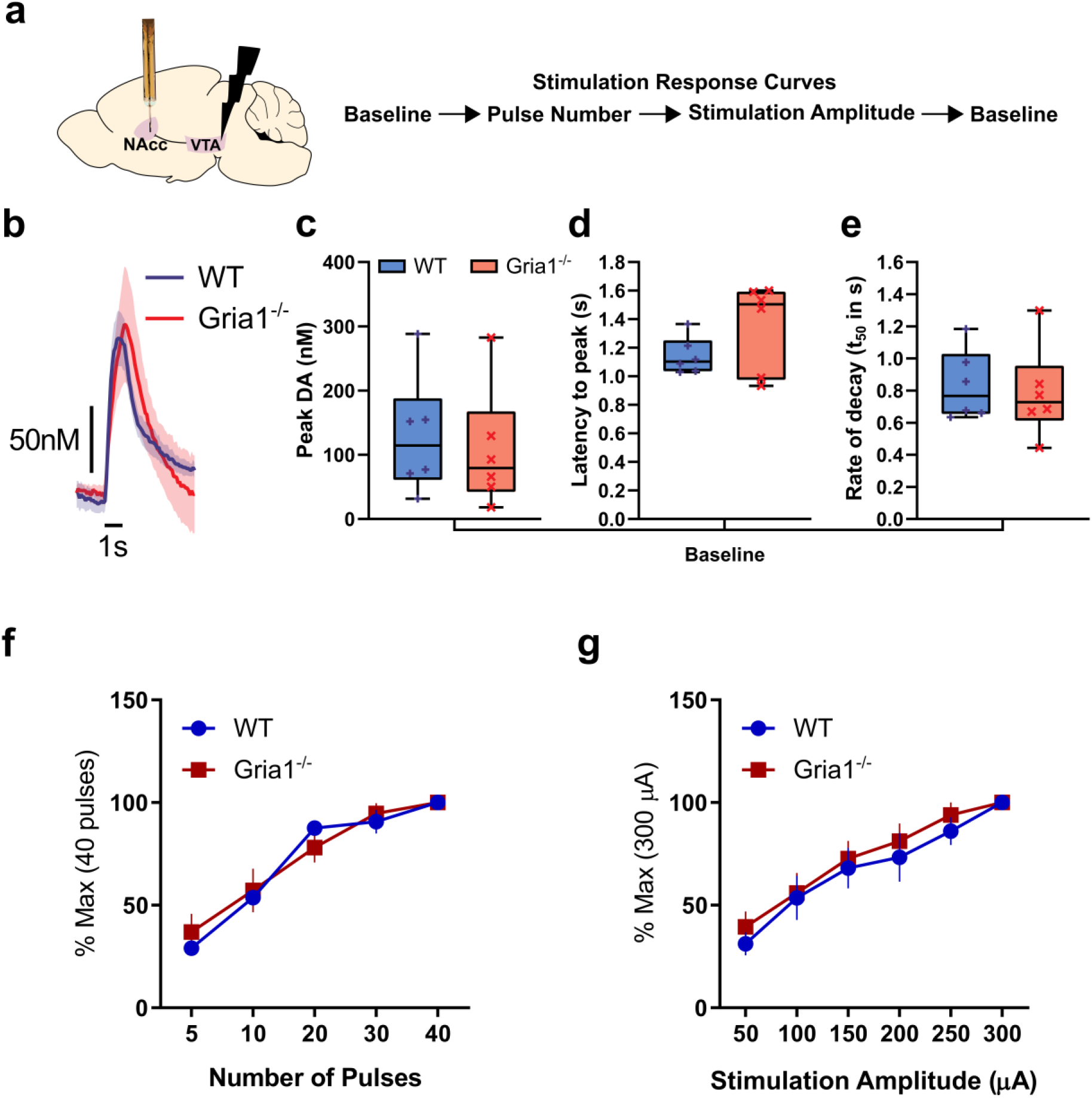
GluA1 deletion does not alter intrinsic VTA-NAcc DA pathway release properties. **(a)** Electrical stimulation of the VTA and FSCV measurement of NAcc DA release in anaesthetized WT and *Gria1*^-/-^ mice. Outline of stimulation protocol involving a baseline session before and after the stimulus response curve protocol. Baseline stimulation parameters were 30Hz, 40 pulses, 2ms pulse width, 300 μA. **(b)** Average traces showing the time course of electrically evoked DA release during baseline stimulations before and after the stimulation response curves (shown in **f** and **g**). Peak DA **(c)**, latency to peak **(d)**, and rate of decay **(e)** of these baseline stimulations did not differ between genotypes. Rate of decay was quantified using the t_50_ from model fit of negative exponential decay for traces during the baseline period with model fit R^2^> 90%. Box and whisker plots show range (error bars), 25^th^-75^th^ percentile (box limits), and median values (line), and individual animal data points. **(f)** Effects of varying number of stimulation pulses on the peak DA release expressed as a percentage of release to the maximum (40) pulses. **(g)** Effects of varying stimulation amplitude on the peak DA release expressed as a percentage of release to the maximum (300 μA) amplitude. Varying the number of pulses and stimulation amplitude significantly changed DA release but this did not depend on genotype. Significant main effect of stimulation amplitude (F_5,45_ = 59.87, *p* < .001), and number of pulses (F_4,45_ = 83.91, *p* < .001), but no significant main effects of genotype or genotype x stimulation interactions (all *Fs* < .59, *p*s > .61, and *Fs* < 1.57, *p*s > .22 respectively). See Supplementary Fig 1 for histology and additional analyses. All error bars represent ± standard error of the mean.

### GluA1 deletion leads to hyper-dopaminergic responses to unsignalled rewards across repeated reward presentations

We next wanted to determine how stimulus-evoked DA in behaving mice was influenced by GluA1 deletion and how this changed with repeated presentation of stimuli. It is well established that reliable NAcc DA responses can be evoked following unpredicted and unsignalled reward delivery [32, 33]. We therefore assessed phasic DA responses to unsignalled rewards (VI120s) in *Gria1*^-/-^ and control mice (see Supplementary Fig 2 for histology).

On average, peak DA responses following reward delivery were markedly higher in *Gria1*^-/-^ than in WT mice (main effect of Genotype: *F*1, 22.48 = 26.45, *p* <.001; Fig 2A). However, closer inspection of the data revealed that this was due to differences in the dynamics of reward-evoked DA across the session in the two groups (Fig 2B). Specifically, while the size of the phasic DA responses decreased progressively across the session in wild-type mice, this decrease was much smaller in *Gria1*^-/-^ mice leading to the development of a hyper-dopaminergic phenotype in these animals. This was supported statistically by a significant Genotype x Reward Number_quadratic_ interaction (*F*_1, 22.20_ = 9.85 *p* = .005). Release was significantly higher in *Gria1*^-/-^ mice after the 6th (*F*_1, 22.69_ = 11.75, *p* = .002), 12th (*F*_1, 22.50_ = 25.74, *p* < .001), 18th (*F*1, 22.50 = 26.36, *p* < .001), and 24th rewards (*F*1, 21.82 = 4.97, *p* = .04). Crucially, however, there were *no* significant differences in reward evoked DA between genotypes to the first reward (*F*_1, 22.87_ = 1.59, *p* = .22).

**Figure 2.**
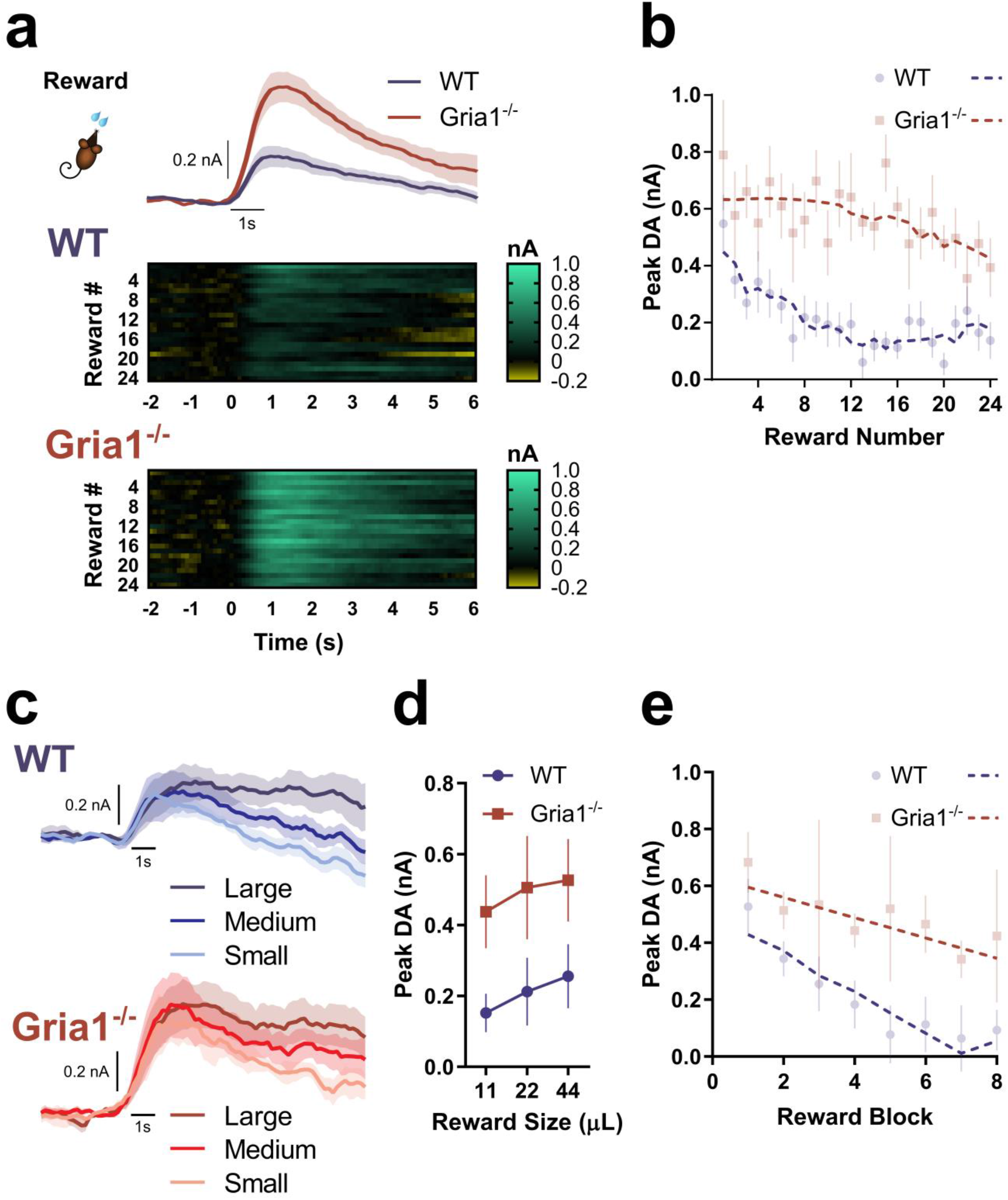
GluA1 deletion leads to hyper-dopaminergic responses to unsignalled rewards as a result of impaired short-term habituation. **(a)** Average DA response to unsignalled reward in WT and *Gria1*^-/-^ mice. Heat plots depict changing DA dynamics within the session over repeated rewards (each reward is a row, and reward is delivered at time = 0). **(b)** Peak DA response plotted as a function of reward number in the session. Average responses depicted as semi-transparent symbols, and predicted values (estimated marginal means) from the statistical model represented as dashed lines. Within-session habituation of reward evoked DA declines rapidly in WT but not *Gria1*^-/-^ mice. **(c)** Average DA response to variable reward sizes (small, medium, large) in WT and *Gria1*^-/-^ mice. A subset of the mice from the previous behavioural task (WT n = 6, KO n = 5 electrodes; WT n = 5, KO n = 4 mice) had patent electrodes for a subsequent test using variable reward sizes (11 μL, 22 μL, 44 μL; small, medium, large rewards respectively). **(d)** Average peak DA response to each reward magnitude. The peak and persistence of the reward evoked DA response were sensitive to reward size (Reward Size *F*_2, 9.00_ = 6.66, *p* = .01; Reward Size x Time *F*_2, 18.22_ = 6.72, *p* = .01). These reward size specific differences in evoked DA did not differ between genotypes (Genotype *F*_1, 9.00_ = 3.56, *p* = .09, Genotype x Reward Size *F*_2, 18.40_ = 0.24, *p* = .79, Genotype x Reward Size x Time *F*_2, 18.22_ = 0.14, *p* = .87). Peak evoked dopamine was significantly greater for large than for small rewards (Sidak corrected threshold of significance *p* < 0.017; Small vs. Medium *F*_1, 18.50_ = 5.22, *p* = .034, Small vs. Large *F*_1, 18.42_ = 13.31, *p* = .002, Medium vs. Large *F*_1, 18.47_ = 1.85, *p* = .19). Reward evoked DA was also significantly higher for large rewards than for medium or small rewards 3s post peak (Small vs. Medium *F*1, 27.41 = 6.01, *p* = .021, Small vs. Large *F*_1, 27.39_ = 27.55, *p* < 0.001, Medium vs. Large *F*1, 27.40 = 7.82, *p* < . 0.01), and 5s post peak (Small vs. Medium *F*1, 21.21 = 4.46, *p* = .047, Small vs. Large *F*1, 21.11 = 23.47, *p* < 0.001, Medium vs. Large *F*1, 21.12 = 7.47, *p* = 0.012). **(e)** Peak DA responses plotted as a function of reward number in the session, in blocks of 3 rewards (averaging across reward size). Average responses depicted as semi-transparent circles, and predicted values (estimated marginal means) from the statistical model represented as dashed lines. All error bars represent ± standard error of the mean.

Next, we tested the sensitivity of the DA responses to changes in reward magnitude (Fig 2C) in a separate recording session. DA signals scaled with reward size in all mice and both genotypes showed an equal sensitivity to reward magnitude (Genotype x Reward Size *F*_2, 18.40_ = 0.24, *p* = .79; Reward Size *F*_2, 18.40_ = 6.66, *p* = .007) (Fig 2D). However, there were again significant genotype differences in the magnitude of the reward response across repeated trials within this session (Genotype x Reward Number x Time *F*_1, 8.99_ = 16.21, *p* = .003) (Fig 2E). Follow up comparisons revealed that while peak DA was again indistinguishable between genotypes at the start of the session, it was significantly greater in *Gria1*^-/-^ than wild-type mice by the last block of rewards (1^st^ Reward: *F*_1, 14.62_ = 0.61, *p* = .45, 4^th^ Reward: *F*_1, 9.13_ = 3.03, *p* = .12, 8^th^ Reward: *F*_1, 14.64_ = 6.07, *p* = .027). Taken together, these findings show that *Gria1*^-/-^ mice exhibit augmented reward-elicited DA release. Importantly, this is an emergent property caused by a progressive decrease in DA responses observed in WTs to repeated presentations of the same reward which is significantly attenuated in *Gria1*^-/-^ mice.

### GluA1 deletion disrupts within-session habituation of light cue evoked DA responses

The data presented above could reflect either a failure in habituation to the sensory-specific stimulus properties of the sucrose reward (e.g., its flavour, texture) or differences in hunger/motivation levels. Striatal DA responses can also be evoked by neutral, non-rewarding stimuli, especially when those stimuli are novel [32, 34–36]. To test whether GluA1 deletion disrupts habituation of the DA response to purely sensory stimuli we next analyzed DA responses to neutral light cues using an established orienting task in which behavioural habituation is sensitive to GluA1 deletion [24, 25].

Presentation of 10 sec neutral light cues evoked robust and reliable phasic DA responses (Fig 3). These signals were comparable in magnitude to the reward-evoked DA (Fig 2), even though these cue-elicited DA responses were independent of any association with reward. Note, we found identical DA responses to a neutral light cue even in naïve mice with no prior exposure to rewards in the operant chamber (Supplementary Fig 5).

**Figure 3.**
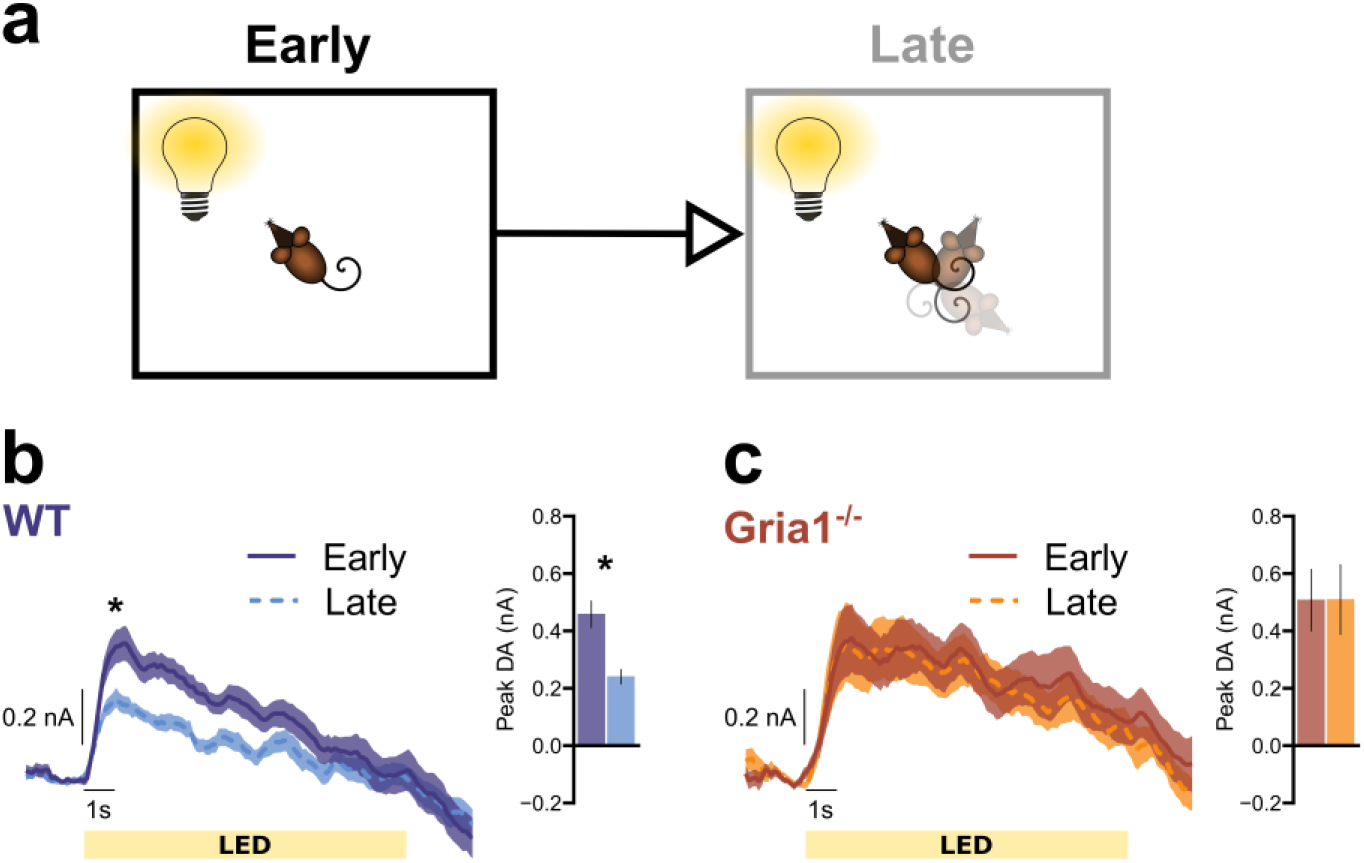
GluA1 deletion disrupts within-session habituation of light cue evoked DA responses. Average DA release to the LED light cue separated into the first (**Early**) and second (**Late**) half of the session to assess stimulus habituation. **(a)** Pictorial representation of relevant task parameters, assessing DA release in response to presentations of the LED light stimulus early and late in the session. **(b)** WT mice show significant habituation of DA responses to the LED. * Simple effects revealed significant differences at 1-4 seconds into stimulus presentation, *F*_1, 15.00_ = 16.29, *p* = .001, *F*_1, 15.50_ = 12.90, *p* = .003, *F*_1, 20.06_ = 6.71, *p* = .017, *F*_1, 24.27_ = 4.38, *p* = .047, respectively. **(c)** *Gria1*^-/-^ mice do not show habituation of DA responses to the LED. Inset bar graphs depict the peak DA estimated at 1s post stimulus onset. All error bars represent ± standard error of the mean.

We first looked for evidence of habituation in evoked DA signals that might occur with repeated presentations of light stimuli across the test session by comparing stimulus elicited DA responses early and late in the session (i.e. first vs second half of the session; Fig 3A). WT mice showed clear evidence of within-session habituation with significantly reduced late session DA signals after repeated presentations of the lights (Fig 3B; WT mice; significant effects of EarlyLate *F*1, 15.00 = 16.29, *p* = .001, EarlyLate x Time *F*_1, 45.48_ = 9.15, *p* = .004). In contrast, DA responses remained high throughout the session in *Gria1*^-/-^ mice with no evidence of habituation of this signal (Fig 3C; *Gria1*^-/-^ mice; no effects of EarlyLate *F*_1, 8.00_ = 0.67, *p* = .44, or EarlyLate x Time *F*_1, 24.24_ = 0.29, *p* = .59 interactions; Genotype differences supported by significant Genotype x EarlyLate interaction *F*_1, 23.00_ = 7.63, *p* = .011, and Genotype x EarlyLate x Time interaction *F*_1, 23.00_ = 6.34, *p* = .019). Importantly, when directly comparing DA release between genotypes, there was significantly higher peak DA in the *Gria1*^-/-^ mice, compared to WT mice, late in the session (*F*_1, 28.60_ = 4.34, *p* = .046), but no significant differences between genotypes early in the session (*F*_1,34.75_ = 0.12, *p* = .727)). Thus, mirroring the earlier results with unsignalled rewards and the robust behavioural impairments in short-term habituation reported previously in *Gria1*^-/-^ mice to a variety of different stimulus types [24, 25], deletion of *Gria1* impairs this within-session habituation of this stimulus elicited DA signal over repeated stimulus presentations, resulting in a hyperdopaminergic phenotype.

### GluA1 deletion disrupts stimulus-specific habituation of light cue evoked DA responses

An important question is whether these observed differences in habituation reflect general changes in arousal and attention in the *Gria1*^-/-^ mice, or are instead a stimulus-specific effect reflecting the recent history of stimulus presentations. We have previously demonstrated *Gria1*^-/-^ deficits in stimulus-specific habituation at the behavioral level in a number of different behavioral settings [25], and replicate this finding in the present task (Figure 4a,b,c). If the habituation of cue evoked striatal DA release is stimulus specific, then the second *“same”* presentation of a target cue in a pairing should elicit lower DA signals than if a *“different”* cue is presented as the second target cue in a pair (e.g. LED→<u>LED</u> DA response < House→<u>LED</u> DA response) (Fig 4a). In WT mice (Fig 4e) presentation of the LED as the second stimulus of a pair elicited less DA if it was preceded by the same stimulus (i.e., the LED) as compared to a different stimulus (house light) (Same vs Different *F*_1, 77.16_ = 4.22, *p* = .043). Thus, WT mice exhibited stimulus-specific habituation of DA responses. In contrast, *Gria1*^-/-^ mice did not show this stimulus-specific habituation effect (Fig 4f); their DA signals to presentation of the second stimulus was the same, irrespective of the identity of the first stimulus (Same vs Different *F*_1, 77.16_ = 0.23, *p* = .627). Interestingly, there was little, if any, stimulus-specific habituation to the house light within a trial in either genotype (Supplementary Fig 3). These Genotype differences were supported by a significant Genotype x HouseLED x Stimulus Novelty interaction (*F*_2, 46.00_ = 3.50, *p* = .038). These stimulus-specific habituation effects in DA responses were also sensitive to the duration of the inter-stimulus interval (Supplementary Figure 4), consistent with theories of habituation and with our previous findings [22]. Thus, we provide evidence that the magnitude of NAcc DA release to unpredicted presentations of sensory stimuli in WT mice undergoes stimulus-specific habituation. Crucially, disruption of this habituation process in *Gria1*^-/-^ mice results in the emergence of a relative amplification of dopaminergic responses to neutral sensory cues.

**Figure 4.**
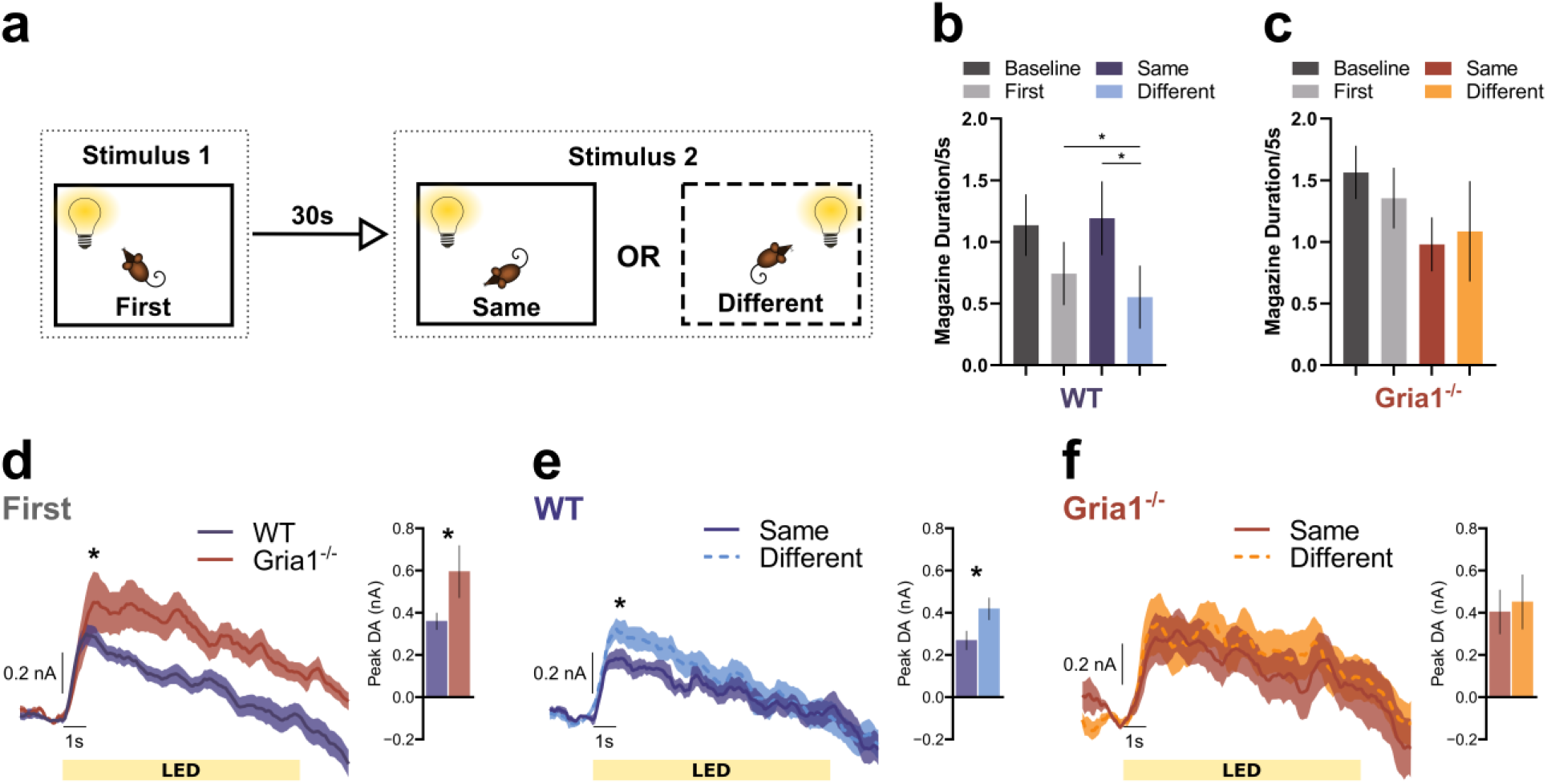
GluA1 deletion disrupts stimulus-specific habituation of light cue evoked dopamine responses. **(A)** Representation of the trial structure in the task. On each trial two stimuli were presented 30s apart. The identity of stimulus 2 in each pair was either the same or different to stimulus 1. Mice have been shown to reduce attention to stimulus 2 when it is the same as stimulus 1, but attention remains high when stimulus 2 is different to stimulus 1 [24, 25]. This selective attentional effect is stimulus specific habituation. Magazine activity in WT **(a)** and KO **(b)** mice immediately prior to stimulus presentation (Baseline), during the first stimulus of a pair (First) and during the second stimulus of a pair depending on whether it is the same or a different stimulus to the first (Same and Different). Suppression of this magazine directed behaviour during stimulus presentation, relative to baseline, provides a measure of a behavioural orienting response [24]. Planned comparisons revealed that WT mice showed significantly less suppression to the Same than the Different stimulus (*F*_1, 44.01_ = 3.18, *p* = .04), and significantly more suppression to the Different stimulus than Baseline (*F*_1, 44.01_ = 4.54, *p* = .04; all remaining *F*_1, 44.01_ < 2.06, *ps* > .16) suggesting that suppression of magazine behaviour to the light stimuli habituated in a stimulus specific manner. In contrast there was no evidence of habituation to the light stimuli in the KO mice (no significant differences between any Periods, all *F*_1, 44.01_ < 3.18, *ps* > .08). This pattern of behaviour replicates earlier findings [24], although the genotype by period interaction did not reach statistical significance (Genotype *F*_1, 14.95_ = 1.44, *p* = .25, Period *F*_3, 44.11_ = 2.08, *p* = .117, Genotype x Period *F*_3, 44.11_ = 1.33, *p* = .28). Average DA release to the LED light stimulus when it was stimulus 1 **(d)**, and when it was stimulus 2 for the WT **(e)** and *Gria1*^-/-^ mice **(f)** . Inset bar graphs depict the peak DA estimated at 1s post stimulus onset. All error bars represent ± standard error of the mean.

## DISCUSSION

Aberrant salience is the dominant mechanism thought to underlie psychosis in neuropsychiatric disorders like schizophrenia, and it is commonly linked to elevated striatal DA [3]. However, there is still little known about its underlying aetiology. Using *Gria1*^-/-^ mice, a mouse model of impaired synaptic plasticity in schizophrenia, here we show the rapid emergence of a hyperdopaminergic state that is directly tied to deficits in short-term habituation in these animals, and thus to inappropriate and maladaptive levels of attention.

Classically, midbrain DA signals and DA release in the NAcc have been attributed to quantitative reward prediction errors that decrease as rewards are effectively predicted by other cues [37–39]. However, there is increasing evidence that the activity of midbrain DA neurons can also be influenced by the sensory properties of stimuli, particularly when those stimuli are novel, salient, and surprising [35, 40–44]. We saw clear, rapid and transient phasic increases in striatal DA levels in response to not only unsignalled naturalistic food rewards but also unsignalled, task irrelevant, neutral light cues. Furthermore, we also provide evidence that a component of these DA responses to unsignalled presentations of both neutral lights and rewards is sensitive to the recent temporal history of stimulus presentations as well as to reward size. This is consistent with recent evidence suggesting that DA neurons may respond more broadly to prediction errors beyond the reward domain [44]. Notably, these DA signals diminished with repeated exposure to the light cues and the rewards in wild-type mice. In contrast, in *Gria1*^-/-^ mice dopamine signals remained high and at levels comparable to the first novel stimulus presentation. Therefore, a hyperdopaminergic state emerged in *Gria1*^-/-^ mice in parallel with the attentional demands of the task. Thus, we can now provide a mechanistic link between the development of a hyperdopaminergic state and aberrant salience in which high levels of attention to stimuli are inappropriately maintained as a result of deficits in habituation.

The present data, combined with previous behavioral studies in *Gria1*^-/-^ mice, demonstrate that GluA1 dysfunction can lead to both aberrant salience and hyperdopaminergic signals in striatum [25]. These phenotypes are highly relevant to psychosis in schizophrenia given the numerous lines of evidence linking GluA1 and associated synaptic plasticity processes with the disorder [11–15] (see also Supplementary Discussion). Furthermore, rather than glutamatergic deficits being distinct from dopaminergic hyperfunction, the present findings show how glutamatergic deficits can directly lead to dopaminergic hyperfunction [2, 45]. Thus, by increasing DA signals to environmental cues and generating inappropriate levels of attention such that stimuli remain “as if novel” when they would otherwise habituate, GluA1 dysfunction effectively increases the window of contiguity which increases the likelihood that associations will form between cues [46, 47]. In this way, GluA1 dysfunction provides a mechanism through which maladaptive associations could be formed between stimuli and events that would otherwise be perceived as unrelated, and thus for generating delusional beliefs.

So, what is the locus of these GluA1 effects? Notably, we found no genotypic differences in electrically evoked DA release in anaesthetized *Gria1*^-/-^ and WT mice, confirming that there were no differences in the intrinsic release properties of DA neurons in the *Gria1*^-/-^ mice. Normal DA release has previously been reported in anaesthetized *Gria1*^-/-^ mice using chronoamperometry in dorsal striatum, although DA clearance was significantly retarded [48]. We found no evidence for altered clearance in the present study and this discrepancy may reflect either regional differences in the effect of GluA1 deletion within the striatum, or differences between the FSCV and chronoamperometry approaches. Nevertheless, these data suggest that the hyperdopaminergic phenotype in *Gria1*^-/-^ mice is unlikely to be due to differences in the intrinsic properties of DA neurons in these mice.

The hippocampus is an attractive alternative candidate. Hippocampal deficits in synaptic function and plasticity, including reduced GluA1, have long been associated with schizophrenia [8, 49–59]. The hippocampus can generate sensory expectations and acts as a comparator to compute surprise signals [60–63]. Consequently, it is implicated in habituation to spatial and non-spatial stimuli [64–67]. A hippocampal surprise signal could then be relayed to DA neurons in the VTA via established polysynaptic circuits [62, 68, 69], thus regulating DA cell activity which, in turn, can promote an increase in attention to novel stimuli via salience networks [70–73] (albeit in a sensory specific manner). A hippocampal surprise signal could also potentially modulate terminal DA release in NAcc directly [74, 75]. Notably, GluA1-dependent plasticity mechanisms in the hippocampus, particularly in CA2/3 subfields [76–78], contribute to the development of short-term memory representations as stimuli become familiar (see Supplementary Discussion). Thus, synaptic plasticity in the hippocampus is likely to be a key regulator of VTA DA cell activity which, in turn, can shape the levels of attention and behavioural resources focused on particular stimuli in the environment. Altered hippocampal-VTA connectivity is a key feature in schizophrenia [79]. Furthermore, habituation to a range of neutral and emotional stimuli is significantly impaired in schizophrenic subjects and this correlates with impaired repetition suppression of BOLD activity in the hippocampus [27, 70, 80–82]. We cannot completely rule out the possibility that this phenotype is a consequence of developmental changes in *Gria1*^-/-^ mice. However, *Gria1*^-/-^ mice do not differ from WT mice on a range of relevant biomarkers [28, 83] and, importantly, the short-term habituation deficit can be rescued in adulthood by reintroduction of GluA1 into the hippocampus of adult *Gria1*^-/-^ mice [78].

Future work is needed to understand how this aberrant salience mechanism might contribute to more complex behaviours with translational relevance to schizophrenia [84–87]. Indeed, Gria1^-/-^ mice show a pattern of deficits consistent with cognitive symptoms in schizophrenia when tested for many of these complex behaviours with touch-screen tasks [25, 88]. This aberrant salience mechanism may account for the mixed efficacy of antipsychotic medication [5, 45, 89], which often targets dopamine D2 receptors. For example, the D2 antagonist haloperidol effectively treats locomotor hyperactivity in *Gria1*^-/-^ mice but, in contrast, has no effect on short-term working memory performance in WT or *Gria1*^-/-^ mice [90]. The mixed efficacy of antipsychotics targeting the dopaminergic system, both in patients and animal models, are again consistent with the idea that striatal hyperdopaminergia and aberrant salience may arise from multiple upstream mechanisms, reflecting the heterogeneity of schizophrenia.

To conclude, striatal hyperdopaminergia and aberrant salience may encompass a range of different mechanisms, reflecting the heterogeneity of schizophrenia. *Gria1* is one of many candidate genes and we do not rule out the possibility that other genetic and environmental disturbances could contribute to aberrant salience via different mechanisms and pathways [2–5, 9, 10]. We also do not exclude the possibility that different molecular mechanisms and different psychological processes underlie the altered DA function reported for dorsal striatum in schizophrenia [27, 91]. Nevertheless, these combined behavioral and neurochemical phenotypes in *Gria1*^-/-^ mice may represent a core mechanism leading to the development of aberrant salience and psychosis as a result of deficits in glutamate synapses and synaptic plasticity in schizophrenia.

## DATA AVAILABILITY

All data and analyses presented in this article are available at: DOI 10.17605/OSF.IO/9F23M

## ACKNOWLEDGMENTS

The work was supported by a Wellcome Trust Senior Research Fellowship to D.M.B (grant no. 087736) and M.E.W. (grant no. 202831/Z/16/Z), Medical Research Council (UK) grant to D.M.B, M.E.W, and P.J.H (grant no. MR/N004396/1), and a BBSRC (UK) Case Studentship to T.B. (grant no. BB/J500446/1). For the purpose of open access, the author has applied a CC BY public copyright license to any Author Accepted Manuscript version arising from this submission. RS receives support from the Ingeborg Ständer Foundation. We would like to thank Greg Daubney for histology, and Katie Hewitt for laboratory facility management and support.

