## Supplementary Materials for "Glutamatergic dysfunction leads to a hyper-dopaminergic phenotype through deficits in short-term habituation: a mechanism for aberrant salience"

**SUPPLEMENTARY INFORMATION**

### Supplementary Methods

#### Electrodes

Carbon fibre electrodes were constructed as described previously [1–3]. Recording electrodes were constructed using 7µm carbon fibre (Goodfellow) encased in a polyamide-coated fused silica capillary tube (outer diameter 90 µm, inner diameter 20 µm; Composite Metal Services Ltd) cut to a length of 0.8-1 cm. One end of the capillary was sealed with epoxy (Devcon), and the exposed carbon fibre protruding from that end was trimmed to a length of 170-200 µm. The other end was glued to s silver pin (Farnell) with silver epoxy (MG Chemicals) and insulated with clear epoxy. Ag/AgCl reference electrodes were constructed using silver wire (Sigma Aldrich) treated with sodium hypochlorite (Fisher Scientific), affixed to a gold pin (Farnell) with silver epoxy and insulated with clear epoxy. Stimulating electrodes were bipolar stainless-steel electrodes (untwisted) measuring 0.15 mm in diameter (PlasticsOne).

#### Anaesthetised recordings

Mice were lightly anaesthetized with isoflurane (3% in oxygen) before receiving an intraperitoneal injection of urethane (25% weight/volume solution; Sigma Aldrich) at a dose of 0.9g/kg, and an i.p. injections of Glycopyrronium bromide (MercuryPharm Ltd) at a dose of 0.01-0.02 mg/kg to minimize adverse bronchial secretions. Initially, anaesthesia was maintained by a combination of the initial urethane dose and isoflurane (0.5-1.5% in oxygen). Isoflurane concentration was gradually lowered during surgery and turned off prior to FCV recordings. Urethane anaesthesia was maintained with top-up injections at 10-20% of the initial dose. Glucose-Saline (0.5% in 0.9% saline; aquapharm) was administered subcutaneously every 3 hours to maintain hydration. Body temperature was monitored and maintained at 35-37C with a rectal probe, heat blanket, and a homeothermic monitor (Harvard Instruments).

Following urethane anaesthesia, mice were placed in a stereotaxic frame (Kopf Instruments) and a local anaesthetic (bupivacaine 2 mg/kg; AstraZeneca) was administered under the scalp. Separate craniotomies were performed to create holes for a recording, stimulating, and a reference electrode, in addition to an anchoring screw (Precision technology Supplies). First an Ag/AgCl reference electrode (anterior-posterior +4.8, medial-lateral +1.0 mm from bregma) and anchoring screw (a-p +2.8, m-l +2.0) were secured to the skull using dental cement (Associated Dental Products Ltd). A carbon fibre recording electrode targeting the NAcc (a-p +1.4, m-l -0.75) was then lowered to -3.40 mm from brain surface and cycled at 60Hz for 20 mins, then cycled at 10Hz for a further 5 mins. A bipolar stimulating electrode, connected to a constant current stimulus isolator (DS-3, Digitimer, Hertfordshire, UK) targeting the VTA (a-p -3.5, m-l +1.0) was lowered to -4.0 mm from brain surface. From these coordinates, the working and stimulating electrodes were progressively lowered (NAcc range -3.5 to -4.75, VTA range -4.0 to -4.80 mm from brain surface) to obtain a maximal evoked DA response. Final depth of NAcc recording electrodes d-v -3.8 ± .2 mm from brain (Supplementary Fig 1A). The VTA was electrically stimulated using the following baseline stimulation parameters: 2ms pulse width, monophasic (+), 30Hz for 40 pulses (1.33s), at an amplitude of 300µA. A minimum of 3 minutes was left between each stimulation when optimizing electrode placements to minimize overstimulation of VTA neurons.

**Fast Scan Cyclic Voltammetry in Freely Moving Animals**

Surgical implantation of electrodes for fast-scan cyclic voltammetry in freely moving animals was performed prior to behavioural pre-training. Mice were anesthetized using isoflurane (4% vol/vol in O2 induction and 1.5% for maintenance) and given buprenorphine (Vetergesic, 0.08 mg/kg) to provide analgesia. Body temperature was maintained at 37 ± 0.5 °C with the use of a homeothermic heating blanket. After induction, the scalp was shaved and cleaned with dilute Hibiscrub (Chlorhexidine Gluconate 4%), 70% alcohol, and a local anesthetic (bupivacaine 2 mg/kg) was injected subcutaneously into the scalp. The skull was then exposed and holes were drilled for the Ag/AgCl reference electrode (a-p -3.1, m-l +1.2 mm from bregma), 4 anchoring screws (Precision Technology Supplies), and for a voltammetric recording electrode in each hemisphere. After the screws and the reference electrode were secured with dental cement, bilateral carbon fiber microelectrodes were lowered into the NAcc (a-p +1.4, m-l ± 1.0 mm from bregma, d-v -3.6 mm from skull). Final depth of NAcc recording electrodes d-v -3.8 ± .2 mm from brain (Supplementary Fig 2A). The carbon fibre and reference electrodes were attached to a 3-pin headstage connector and secured with dental cement. Following surgery, animals were again administered an analgesic (buprenorphine 0.08 mg/kg) and a non-steroidal anti-inflammatory drug (NSAID; Metacam, 5 mg/kg), and given palatable food to facilitate post-operative recovery. NSAIDs were also administered for at least 3 d following surgery. Mice were left for at least 2 weeks post-operative recovery before behavioural recording.

**Fast-scan cyclic voltammetry**

Fast-scan cyclic voltammetry recordings were performed as described previously [2–5]. In brief, voltammetric scans were performed at a frequency of 10 Hz throughout the session. Prior to a scan, the carbon fiber was held at a potential of −0.4 V (versus Ag/AgCl) and then, during the scan, ramped up to +1.3 V and back to −0.4 V at 400 V/s. The application of this waveform causes redox reactions in electrochemically active species, such as dopamine, at the surface of the carbon fiber, which can be recorded as changes in current over time. Based on previously established criteria, the recorded current to un-cued delivery of sucrose reward (20%) obtained at the start and end of the session was used to verify the chemical sensitivity of the recording electrode to dopamine on a given session. An extracted cyclic voltammogram was linearly regressed against a dopamine standard, with $R^2 \geq 0.75$ set as the criterion based on the discriminability of dopamine from other common neurochemicals in a flow cell. Only sessions where sufficient discriminability was confirmed were included in the analysis presented.

**Behavioural apparatus**

Mice were tested in operant chambers (15.9 x 14.0 x 12.7 cm; ENV-307A, Med Associates), enclosed in sound attenuating cubicles (ENV-022MD, Med Associates), using scripts programmed with Med-PC IV software (Med Associates). A custom-made gravity-based reward-delivery system was used to deliver 20 % sucrose solution (22 µL) into a custom-built food magazine via a spout. The food magazine extended out into the box, as opposed to the standard food magazine used in previous studies which was recessed into the wall of the operant box [6]. This modification was made to allow head-capped/tethered mice clear access to the reward in the food magazine. One consequence of this was that mice didn't need to withdraw their heads from the magazine to see the light stimuli. Thus, we found that measurement of orienting to light stimuli, in terms of suppression of magazine activity, was not as robust as previously reported [6]. Two LED (ENV-321M, Med Associates) lights were positioned on the same wall as the food magazine to the left and to the right of the magazine. A house light (ENV-315M, Med Associates) was placed on top of the conditioning box facing away from the chamber floor. Each chamber was equipped with a fan (ENV-025AC, Med Associates) that was turned on for the duration of the session.

**Calibration of electrodes for Anaesthetised recordings**

Electrodes were cycled at 60Hz for 5-10 mins in flowing artificial cerebrospinal fluid (ACSF; 154.7 mM $Na^+$, 0.82 mM $Mg^{2+}$, 2.9mM $K^+$, 132.49 mM $CL^-$, 1.1 mM $Ca^{2+}$ in deionized water, brought to pH ~7.4 with hydrochloric acid; Sigma Aldrich). ACSF was switched to 1000 nM dopamine hydrochloride solution (mixed in ACSF; Sigma Aldrich) for 30 seconds. Signals to the dopamine solution increased but did not reach a stable plateau, so the inflection point (the point at which the rate of increase plateaued) was used in lieu of a maximum. The failure to reach a maximum and continued increase in signal size past the inflection point is likely due to adherence of dopamine to the electrode surface [3].

**Voltammetric Analysis**

Voltammetric analysis was initially carried out using software written in LabVIEW (National Instruments). Data were baselined 0.5s prior to the event of interest and low-pass filtered at 2 kHz before chemometric analysis was performed using principal components regression [7, 8] on custom functions written in Matlab. As described previously [2], a principal components analysis using a standard training set of stimulated dopamine release detected by chronically implanted electrodes was used to discriminate between changes in dopamine concentration from other unrelated electrochemical fluctuations such as pH changes. Retained principal components are then regressed against test data using an inverse least squares regression. Any trial data that did not sufficiently fit the principal components regression model (model rejected on > 20% of data points) were excluded from analysis. In anaesthetised recordings, electrodes were pre-calibrated using a flow cell to establish a conversion between current signals (nA) and dopamine concentration (nM).

**Exclusion criteria**

In experiments with FSCV recordings in freely moving animals, 40 male mice (WT n = 22, *Gria1*$^{-/-}$ n = 18) were used in total, including for method development, piloting of electrode placements and optimization of dopamine recordings. DA signals were required to meet an established criteria of DA detection ($R^2 \geq 0.75$ between a dopamine standard and an extracted cyclic voltammogram in response to pre- or post-session reward [1, 2]).4 mice were excluded following histological assessment of brain tissue due to misplaced electrodes (WT n = 1, *Gria1*$^{-/-}$ N = 3) outside the NAcc. A final cohort of N=14 male mice (WT n=9, Gria1-/- n=5) with a total of N = 25 working electrodes (WT n=16, *Gria1*$^{-/-}$ n=9 electrodes), located in the nucleus accumbens core (NAcc), contributed to the dataset reported in this study.

**Statistical Analysis**

Further statistical analysis was performed using SPSS v25 (IBM). For anaesthetised recordings, peak DA levels were analyzed using a mixed ANOVA with between subjects factors of Genotype (WT, KO) and sex (Male, Female). Analysis of the stimulation amplitude response curve also contained a within-subjects factor of stimulation amplitude (50, 100, 150, 200, 250, 300 µA). Analysis of the pulse number response curve also contained a within-subjects factor of pulse number (5, 10, 20, 30, 40 pulses).

In freely moving voltammetry recordings, continuous light evoked FSCV traces were analyzed using a linear mixed effects model. Genotype (WT, KO), HouseLED (House light, LED stimulus identity), and Stimulus (first light, same, different) were defined as categorical fixed factors, and Time was defined as a continuous fixed factor sampled at 10Hz. Time was measured from 1s post stimulus onset to stimulus offset and centered at 1s. This allowed for a linear signal (signals peaked at ~1s post stimulus onset) and for meaningful interpretations of interaction terms at the peak of the signal. Accordingly, any effects that did not interact with Time represent differences between peak DA release, and any effects that did interact with Time represent differences in the linear slopes of the signals. Reward evoked peak DA responses in these sessions were also analyzed using linear mixed effects models. Fixed factors were Genotype (WT, KO), and reward number (up to 24 rewards). Reward number was centered and an additional reward number x reward number fixed factor was entered in the model to account for any quadratic curvature in the data. Subject was defined as each mouse x electrode x session. Note however the pattern of results remained the same when considering only mouse or mouse x electrode as subjects.

A similar analysis was conducted for the separate test session involving variable reward sizes. Reward evoked DA was taken from 1s post reward delivery, and the subsequent 5s of DA signals were analysed. This analysis included main factors of Genotype, Reward size (small, medium, large), Reward number (1-8), and time (5s of data sampled at 10Hz). Again, reward number was centered to allow for a quadratic component, however this was removed from the final analysis as there did not appear to be any curvature in the data.

All analyses were conducted using the MIXED command in SPSS v.25, and a variance components (VC) G matrix was specified as part of the RANDOM subcommand, and used a restricted maximum likelihood estimation of effect parameters. Random factors included an intercept and the full factorial structure of all the repeated factors. Denominator degrees of freedom in F tests were calculated using the Satterthwaite approximation which is valid for unbalanced experimental designs. Any situations where reward delivery and stimulus presentation overlapped were excluded from all analyses.

In freely moving voltammetry recordings, summary plots of peak DA signals were calculated from the linear mixed effects model by estimating the marginal means using both fixed and random effects parameter estimates. This generated a predicted peak DA value for each subject level at each experimental condition. Note that these model predictions of peak DA were similar to raw signals at the signal peak (~1s post stimulus or reward onset) reflecting an appropriate model fit.

Magazine behaviour from the freely moving voltammetry recordings was first processed to remove any trials in which the baseline responding was missing for at least 10s prior to the first cue as this baseline is critical to interpreting any cue elicited suppression in responding. Duration of time (s) spent in the magazine during the first 5s of each 10s light stimulus was analyzed as this has been shown to be most sensitive to attentional orienting responses [9]. Inspection of the full 10s of each cue revealed a similar pattern of results. A Mixed Model ANOVA (SPSS MIXED command) was conducted with factors of Genotype (WT, KO) and a repeated factor of Period (Baseline, First, Same, Different), and a random factor of Subject, using a compound symmetry covariance matrix (best model fit using Schwarz’s BIC).

### Supplementary Results

**GluA1 deletion does not alter sensitivity to electrically evoked VTA-NAcc DA release in anaesthetized mice**

Electrically evoked VTA-NAcc dopamine levels did not differ between genotypes at a range of stimulation amplitudes and pulse numbers (Fig 1). These data were presented as a percentage of the maximum stimulation parameter (Fig 1). The raw signals reveal the same pattern of results (Supplementary Fig 1) and statistical significances and are presented below for completeness.

There was no evidence of a genotype difference in electrically stimulated VTA-NAcc dopamine levels. This was true for 5, 10, 20, 30, and 40 pulses (Supplementary Fig 1B ; significant main effect of Pulses $F_{4,45} = 11.53$, $p < .001$, but no main effect or interaction with genotype, $F_{1,9} = 0.23$, $p = .64$, $F_{4,45} = 0.34$, $p = .85$ respectively) and 50, 100, 150, 200, 250, and 300 µA stimulation amplitudes (Supplementary Fig 1C; significant main effect of stimulation amplitude $F_{5,45} = 9.71$, $p < .001$, but no main effect or interaction with genotype, $F_{1,9} = 0.02$, $p = .91$, $F_{5,45} = 0.10$, $p = .99$ respectively), demonstrating that GluA1 deletion does not lead to intrinsic differences in release sensitivity or the balance of DA release/reuptake in this subcortical dopamine pathway.

Surprisingly, there was a significant main effect of sex ($F_{1,9} = 10.30$, $p = .01$) and interaction with Pulses ($F_{5,45} = 2.79$, $p = .03$), but no interactions with Genotype (all *F*s < .04, *p*s > .89; Supplementary Fig 1B). Similarly, there was a significant main effect of sex ($F_{1,9} = 11.78$, $p = .007$) and interaction with stimulation amplitude ($F_{5,45} = 3.29$, $p = .02$), but no interactions with Genotype (all *F*s < .21, *p*s > .93 Supplementary Fig 1C). Closer inspection revealed that stimulated DA release was lower in female than male mice (Supplementary Fig 1d-e). While it is unclear why we observe such large sex differences, however they are consistent with the presence of sex differences in reward pathways[10]. Importantly, sex did not interact with Genotype.

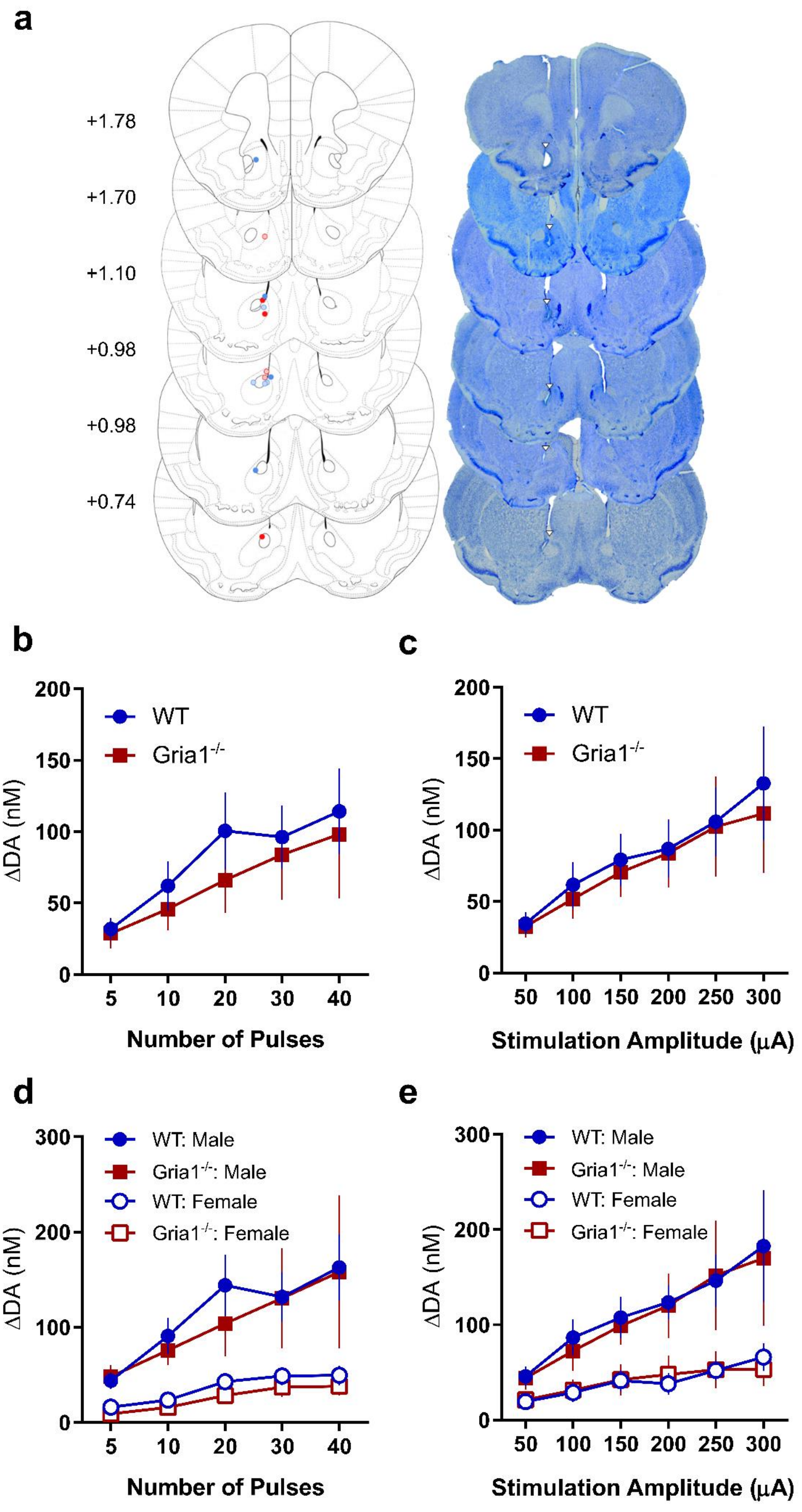
a
+1.78
+1.70
+1.10
+0.98
+0.98
+0.74
b
200
150
100
50
0
WT
Gria1-/-
ΔDA (nM)
5
10
20
30
40
Number of Pulses
c
WT
Gria1-/-
ΔDA (nM)
50
100
150
200
250
300
Stimulation Amplitude (μA)
d
300
WT: Male
Gria1-/-: Male
WT: Female
Gria1-/-: Female
ΔDA (nM)
Number of Pulses
e
WT: Male
Gria1-/-: Male
WT: Female
Gria1-/-: Female
ΔDA (nM)
Stimulation Amplitude (μA)

**Supplementary Figure 1. GluA1 deletion does not alter intrinsic VTA-NAcc DA pathway release properties**

**(a)** Representative location of working electrodes in NAcc and in anaesthetised experiments. Placements are depicted (Left) with WT (blue circles) and Gria1$^{-/-}$ (red circles) separated by male (semi-transparent circles) and female (opaque circles). Representative samples (Right) of Nissl stained sections showing electrode lesion sites (white triangle) in different mice at each anterior-posterior coronal section (numbers indicate mm relative to bregma). **(b)** Effects of varying number of stimulation pulses on the peak DA release (nM). **(c)** Effects of varying stimulation amplitude on the peak DA release (nM). These raw data are also presented as a percentage of the maximum stimulation in Figure 1f-g. Male and female mice in each genotype are plotted separately for varying **(d)** number of pulses and **(e)** stimulation amplitude. All error bars represent ± standard error of the mean.

**GluA1 deletion leads to hyper-dopaminergic responses to unsignalled rewards which develops with repeated reward presentation**

DA responses to unsignalled rewards were assessed in a 48 min session with sucrose rewards (22 µL) delivered contingent upon an operant nose poke response on average every 2 mins. The following analyses provide a complete description of the results described in the main text. Peak DA response to rewards in the session was significantly higher in *Gria1*$^{-/-}$ than in WT mice (Fig 2A). However, this was due to significantly impaired within-session habituation to the reward throughout the session in the knockout mice (Fig 2B). This observation was supported statistically by a significant main effect of Genotype ($F_{1, 22.48}$ = 26.45, $p$ < .001), and a significant Genotype x Reward Number$_{quadratic}$ interaction ($F_{1, 22.20}$ = 9.85 $p$ = .005; significant main effect of Reward Number $F_{1, 22.20}$ = 14.15, $p$ = .001, but no main effect of Reward Number$_{quadratic}$ $F_{1, 22.20}$ = 1.60, $p$ = .22, or Genotype x Reward Number interaction $F_{1, 22.65}$ = 0.07, $p$ = .80). While there was no significant differences between genotypes following the first reward ($F_{1, 22.87}$ = 1.59, $p$ = .22), reward evoked DA was significantly higher in *Gria1*$^{-/-}$ mice after the 6th ($F_{1, 22.69}$ = 11.75, $p$ = .002), 12th ($F_{1, 22.50}$ = 25.74, $p$ < .001), 18th rewards ($F_{1, 22.50}$ = 26.36, $p$ < .001), and at 24th reward ($F_{1, 21.82}$ = 4.97, $p$ = .04).

Next, we tested the sensitivity of the reward evoked DA response in *Gria1*$^{-/-}$ mice to changes in reward magnitude (Fig 2C). A subset of the mice from the previous behavioural task (WT n = 6, *Gria1*$^{-/-}$ n = 5 electrodes; WT n = 5, *Gria1*$^{-/-}$ n = 4 mice) had patent electrodes for a subsequent test using variable reward sizes (11 µL, 22 µL, 44 µL; small, medium, large rewards respectively). Total evoked DA signals were sensitive to reward size in all mice (Fig 2D). These differences were present both in the peak and persistence of the evoked DA response (Reward Size $F_{2, 9.00}$ = 6.66, $p$ = .01; Reward Size x Time $F_{2, 18.22}$ = 6.72, $p$ = .01). These reward size specific differences in evoked DA did not differ between genotypes (Genotype $F_{1, 9.00}$ = 3.56, $p$ = .09, Genotype x Reward Size $F_{2, 18.40}$ = 0.24, $p$ = .79, Genotype x Reward Size x Time $F_{2, 18.22}$ = 0.14, $p$ = .87). Peak evoked dopamine was significantly greater for large than for small rewards (Sidak corrected threshold of significance $p < 0.017$; Small vs. Medium $F_{1, 18.50} = 5.22$, $p = .034$, Small vs. Large $F_{1, 18.42} = 13.31$, $p = .002$, Medium vs. Large $F_{1, 18.47} = 1.85$, $p = .19$). Reward evoked DA was also significantly higher for large rewards than for medium or small rewards 3s post peak (Small vs. Medium $F_{1, 27.41} = 6.01$, $p = .021$, Small vs. Large $F_{1, 27.39} = 27.55$, $p < 0.001$, Medium vs. Large $F_{1, 27.40} = 7.82$, $p < . 0.01$), and 5s post peak (Small vs. Medium $F_{1, 21.21} = 4.46$, $p = .047$, Small vs. Large $F_{1, 21.11} = 23.47$, $p < 0.001$, Medium vs. Large $F_{1, 21.12} = 7.47$, $p = .012$). Therefore, *Gria1*$^{-/-}$ and WT mice were equally sensitive to changes in reward size.

While there were no apparent differences between genotypes in sensitivity to reward magnitude, there were significant genotype differences in the habituation of these reward evoked signals across the session (significant effects of Reward Number $F_{1, 8.93} = 18.30$, $p = .002$, Genotype x Reward Number x Time $F_{1, 8.99} = 16.21$, $p = .003$, but no significant effects of Genotype x Time $F_{1, 8.99} = 0.01$, $p = .92$,Genotype x Reward Number $F_{1, 8.93} = 2.72$, $p = .13$, Reward Number x Time $F_{1, 8.99} = 3.26$, $p = .11$, Genotype x Reward Size x Reward Number $F_{2, 18.14} = 0.50$, $p = .61$, Genotype x Reward Size x Reward Number x Time $F_{2, 17.65} = 0.51$, $p = .61$; Fig 2E). Follow up comparisons revealed that peak DA was significantly greater in *Gria1*$^{-/-}$ than WT mice on the last block of rewards in the session (Reward Number 1 $F_{1, 14.62} = 0.61$, $p = .45$, Reward Number 4 $F_{1, 9.13} = 3.03$, $p = .12$, Reward Number 8 $F_{1, 14.64} = 6.07$, $p = .027$). This suggests that peak reward evoked DA responses in *Gria1*$^{-/-}$ mice did not habituate within-session to the same extent as WT mice. Notably, this genotype difference in within-session habituation was not present 5s post peak (Reward Number 1 $F_{1, 24.78} = 2.47$, $p = .13$, Reward Number 4 $F_{1, 17.52} = 1.55$, $p = .23$, Reward Number 8 $F_{1, 24.79} = 0.28$, $p = .60$). These findings successfully replicate the deficit of within-session habituation of the reward response in the *Gria1*$^{-/-}$ mice.

***Gria1*$^{-/-}$ disrupts within-session habituation of light cue evoked DA response**

Analysis of both House and LED lights simultaneously is presented here. This analysis revealed that the WT mice showed elevated DA release to the House light compared to the LED (comparison of House and LED in WT mice $F_{1, 23.00} = 19.29$, $p < .001$; Fig 3; Supplementary Fig 2B-D), whereas this stimulus identity specific effect was not evident in the *Gria1*$^{-/-}$ mice (comparison of House and LED in *Gria1*$^{-/-}$ mice $F_{1, 23.00} = 0.00$ $p = .95$; Genotype x HouseLED interaction $F_{1, 23.00} = 7.21$, $p = .013$). Notably, this was not simply due to differences in within-session habituation effects in the WT mice (no significant interactions between HouseLED x EarlyLate $F_{1, 23.01} = 3.12$, $p = .019$, Genotype x HouseLED x EarlyLate $F_{1, 23.01} = 0.56$, $p = .46$, Genotype x HouseLED x EarlyLate x Time $F_{1, 23.01} = 0.53$, $p = .47$). It is possible that these stimulus specific differences are due to intrinsic differences in the salience of the House and LED lights. The LED provides a punctate local stimulus whereas the House light is a more diffuse cue which might be more akin to an occasion setter or contextual cue. In support of this, the houselight in the present study was also located significantly further from the floor of the operant chamber (unlike previous reports [6]) to accommodate a commutator for voltametric recording, and this may account for differences in stimulus processing between the House light and LED.

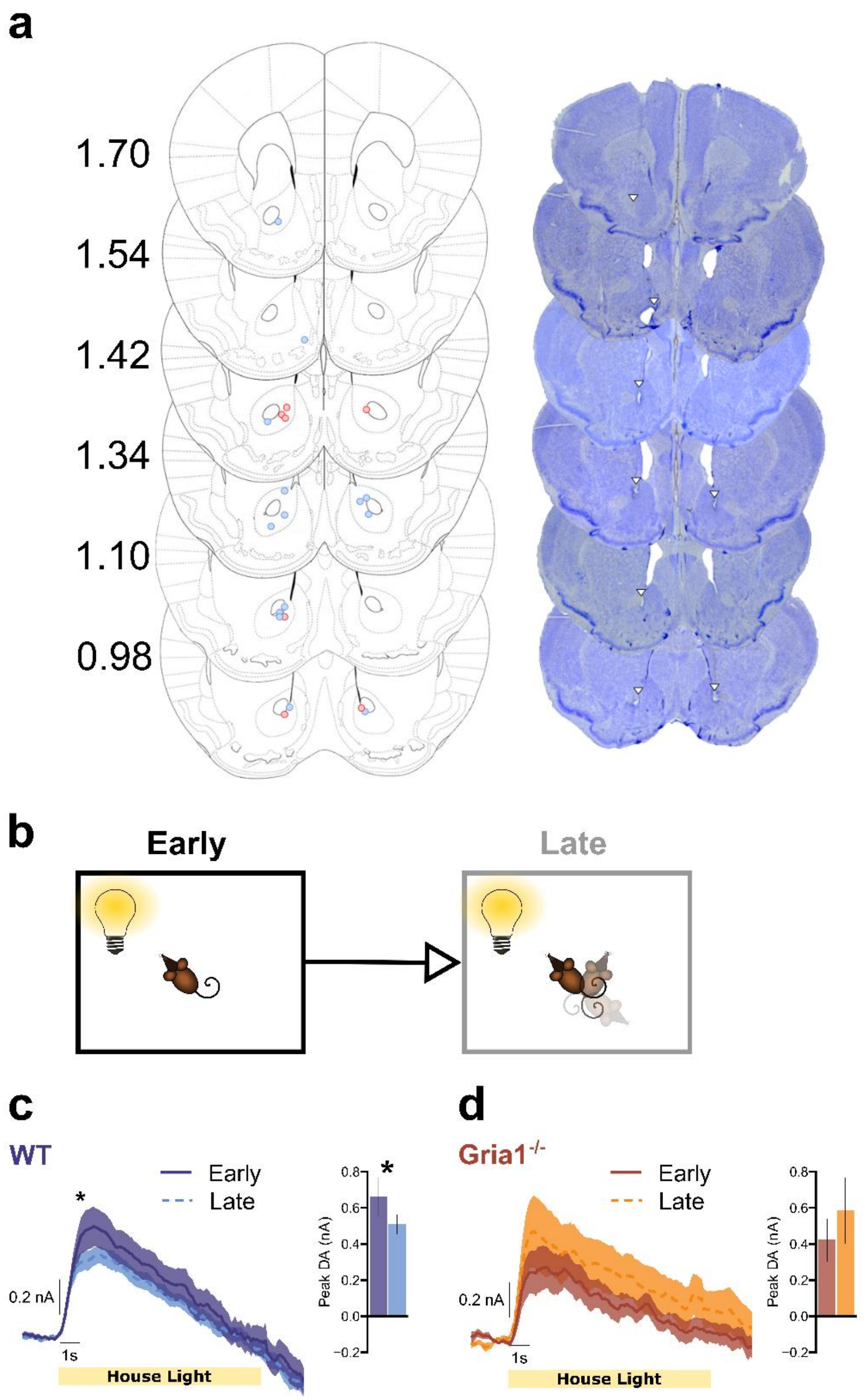


**Supplementary Figure 2. GluA1 deletion disrupts within-session habituation of light cue evoked DA responses**

Representative location of working electrodes in NAcc in behavioural experiments. Average DA release to the House light cue separated into the first (**early**) and second (**late**) half of the session to assess stimulus habituation. **(a)** Placements are depicted (Left) with WT (blue circles) and Gria1$^{-/-}$ (red circles). Representative samples (Right) of Nissl stained sections showing electrode lesion sites (white triangle) in different mice at each anterior-posterior coronal section (numbers indicate mm relative to bregma). **(b)** Pictorial representation of relevant task parameters, assessing DA release in response to presentations of the LED light stimulus early and late in the session. DA responses to the House light reflecting significant within-session habituation in WT **(c)** but not in *Gria1*$^{-/-}$ **(d)** mice. All error bars represent ± standard error of the mean.

#### *Gria1*$^{-/-}$ disrupts stimulus-specific habituation of light cue evoked dopamine

Here we analyse the stimulus specificity of habituation in stimulus evoked striatal DA release extending the analysis in the main text (Fig 4d-f) to include both the House light and the LED light (Supplementary Fig 3). If cue evoked striatal DA release is stimulus specific, then the same cue presented twice should elicit lower DA than if a different cue is presented (i.e. stimulus-specific habituation). WT mice exhibited this stimulus specific habituation of the DA response (Same vs Different $F_{1, 77.16} = 4.22$, $p = .043$) when the target cue was an LED light, but not the House light (no significant differences in House light for Same vs Different $F_{1, 77.16} = 0.84$, $p = .36$; significant HouseLED x Stimulus Novelty interaction in WT mice $F_{2, 30.01} = 5.00$, $p = .013$; Fig4, Supplementary Figure 3C). In contrast, *Gria1*$^{-/-}$ mice did not show differences in responding to the LED or House light regardless of whether it was the same or different to the first light in the trial (no significant effect in *Gria1*$^{-/-}$ mice of HouseLED $F_{1, 8.00} = 0.29$, $p = .61$, Stimulus Novelty $F_{2, 16.00} = 0.50$, $p = .62$, or HouseLED x Stimulus Novelty interaction Stimulus $F_{2, 16.00} = 2.13$, $p = .15$). This suggests that the stimulus specific habituation observed in the WT mice was sensitive to the counterbalanced stimulus identity (main effect of HouseLED $F_{1, 23.00} = 7.96$, $p = .01$, Time ($F_{1, 23.00} = 74.11$, $p < .001$), and a HouseLED x Time interaction $F_{1, 23.00} = 24.49$, $p < .001$; All remaining effects $F$s < 3.02, $p$s > .06). However this effect was abolished in the *Gria1*$^{-/-}$ mice. These Genotype differences were supported by a significant Genotype x HouseLED x Stimulus Novelty interaction ($F_{2, 46.00} = 3.50$, $p = .038$).

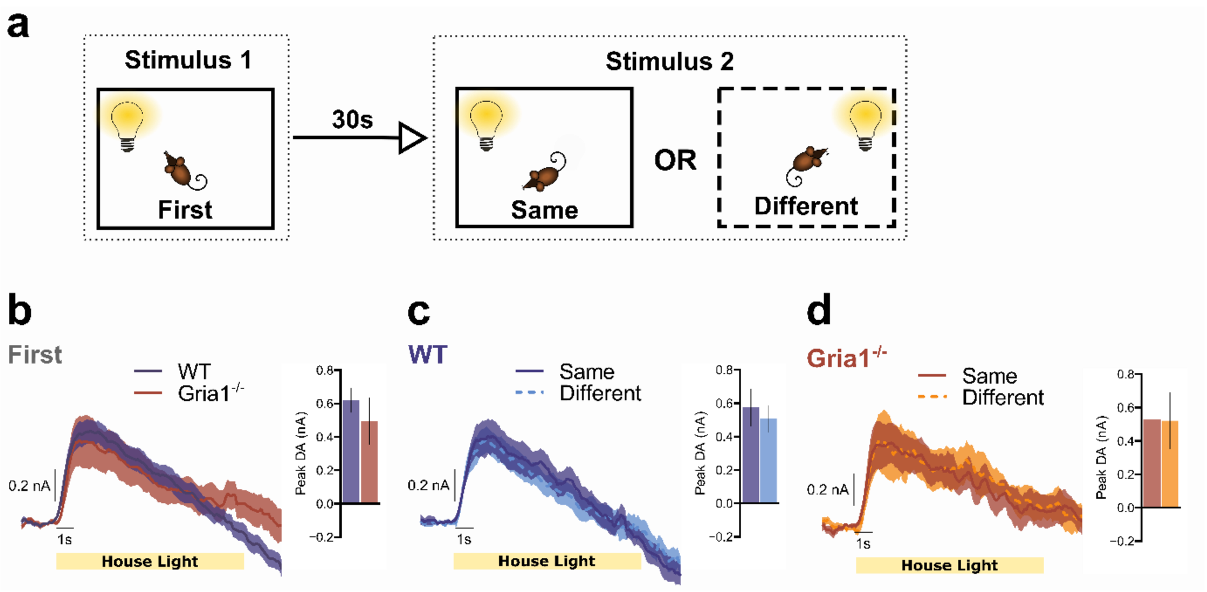


**Supplementary Figure 3. GluA1 deletion disrupts stimulus-specific habituation of light cue evoked dopamine responses**

**(a)** Representation of the trial structure in the task. On each trial two stimuli were presented 30s apart. The identity of stimulus 2 in each pair was either the same or different to stimulus 1. These data correspond to those presented in Figure 4, but now showing responses to the House light. Average DA release to the House light stimulus when it was stimulus 1 **(b)**, and when it was stimulus 2 for the WT **(c)** and *Gria1*$^{-/-}$ mice **(d)**. All error bars represent ± standard error of the mean.

***Do the House and LED lights have different arousing properties?***

Differences in the DA responses to the target (second) cue could not simply be explained in terms of differences in the arousing properties of the House Light and the LED when they were the first stimulus in a pairing. Importantly, there were no differences between House→House and LED→House trials (WT $F_{1,\,48.08} = 2.42$, $p = .127$, *Gria1*$^{-/-}$ $F_{1,\,28.77} = 0.07$, $p = .796$).

**Are there differences in DA release to the House and LED lights?**

*Gria1*$^{-/-}$ mice exhibited significantly greater DA release than WT mice to the first stimulus when it was an LED when averaged across the whole session ($F_{1,\,44.00} = 4.66$, $p = .036$, Fig 4d); but not for the House light ($F_{1,\,37.23} = 0.59$, $p = .45$, Supplementary Fig 3b). This difference for the LED is likely to reflect the significant within-session habituation to the LED in the WT mice, and a lack of within-session habituation in the *Gria1*$^{-/-}$ mice (as reported in Fig 3).

Similar to the previous analysis of within-session habituation, we found significant stimulus identity differences in the sensitivity of the evoked DA response to the House light and LED between the WT and *Gria1*$^{-/-}$ mice (significant main effect of HouseLED $F_{1,\ 23.00}$ = 7.96, $p$ = .01, HouseLED x Time interaction $F_{1,\ 23.00}$ = 26.49, $p$ < .001, Genotype x HouseLED interaction $F_{1,\ 23.00}$ = 4.49, $p$ = .045, but no effect of Genotype x HouseLED x Time $F_{1,\ 23.00}$ = 2.88, $p$ = .10). WT mice had significantly greater evoked DA to the House light than to the LED ($F_{1,\ 23.00}$ = 16.95, $p$ < .001), whereas *Gria1*$^{-/-}$ mice exhibited similar levels of peak DA to both stimuli ($F_{1,\ 23.00}$ = 0.19, $p$ = .67).

**GluA1 deletion disrupts stimulus-specific habituation of behavioural orienting responses**

Figure 4 depicts magazine activity in WT **(Figure 4b)** and KO **(Figure 4c)** mice immediately prior to stimulus presentation (Baseline), during the first stimulus of a pair (First) and during the second stimulus of a pair depending on whether it is the same or a different stimulus to the first (Same and Different). Suppression of this magazine directed behaviour during stimulus presentation, relative to baseline, provides a measure of a behavioural orienting response [6]. Planned comparisons revealed that WT mice showed significantly less suppression to the Same than the Different stimulus ($F_{1,\ 44.01}$ = 3.18, $p$ = .04), and significantly more suppression to the Different stimulus than Baseline ($F_{1,\ 44.01}$ = 4.54, $p$ = .04; all remaining $F_{1,\ 44.01}$ < 2.06, *ps* > .16), suggesting that suppression of magazine behaviour to the light stimuli habituated in a stimulus specific manner. In contrast there was no evidence of habituation to the light stimuli in the KO mice (no significant differences between any Periods, all $F_{1,\ 44.01}$ < 3.18, *ps* > .08). This pattern of behaviour replicates earlier findings [6], although the genotype by period interaction did not reach statistical significance (Genotype $F_{1,\ 14.95}$ = 1.44, $p$ = .25, Period $F_{3,\ 44.11}$ = 2.08, $p$ = .117, Genotype x Period $F_{3,\ 44.11}$ = 1.33, $p$ = .28). This likely reflects the use of a modified magazine receptacle that was not enclosed in order to allow implanted mice to access sucrose. Therefore, in contrast to earlier studies with enclosed magazines, a mouse could see the light stimuli without needing to disengage completely from the magazine area, thus potentially reducing the sensitivity of the behavioural measure. However, crucially the overall pattern of results replicates earlier behavioural findings, and matches the stimulus sensitivity of the measured DA responses.

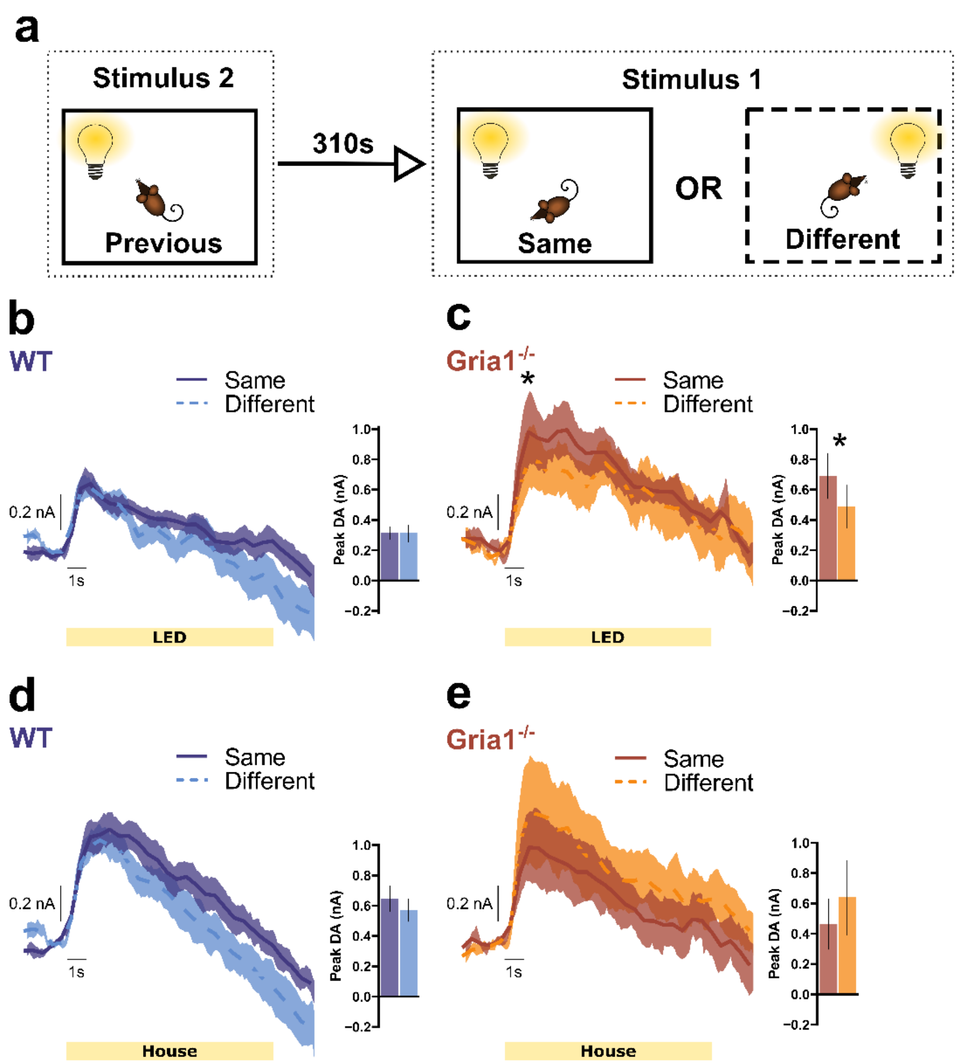


**Supplementary Figure 4. Stimulus-specific habituation of cue evoked dopamine in WT mice is not present after a 310s inter-stimulus-interval.**

**(a)** Representation of the trial structure analysed to assess the effects of a longer inter-stimulus interval (310 sec). The identity of Stimulus 1 (trial n) was separated by whether it was the same or different to Stimulus 2 (trial n-1), presented 310s earlier, at the end of the previous trial. This analysis is comparable to Figure 4 except that the inter-stimulus-interval is now 310s instead of 30s. Stimulus specific habituation is sensitive to the time interval between stimulus presentations and is weaker at longer time intervals. Average DA release to Stimulus 1 (trial n) when it was the LED light in WT **(b),** and *Gria1*$^{-/-}$ mice **(c),** and when it was the house light in WT **(d),** and *Gria1*$^{-/-}$ mice **(e).** Inset bar graphs depict the peak DA estimated at 1s post stimulus onset. All error bars represent ± standard error of the mean.

In WT mice, there was now no evidence of stimulus-specific habituation of the peak DA response after 310 s for either the LED (WT LED: Same vs Different, $F_{1, 32.35} = 0.04$, $p = .851$) or the house light (WT House: Same vs Different, $F_{1, 32.35} = 2.22$, $p = .146$), suggesting dishabituation of the response after this longer interval. Peak DA in *Gria1*$^{-/-}$ mice was significantly higher to the LED in the same condition after a 310 s interval (*Gria1*$^{-/-}$ LED: Same vs Different, $F_{1, 32.20} = 5.58$, $p = .024$), but not for the house light (*Gria1*$^{-/-}$ House: Same vs Different, $F_{1, 32.20} = 4.13$, $p = .050$). Furthermore, peak DA to the same LED in *Gria1*$^{-/-}$ mice was significantly higher than to same LED in WT mice (LED Same: WT vs KO $F_{1, 53.59} = 5.86$, $p = .019$), but did not differ for the different LED (LED Different: WT vs KO, $F_{1, 55.03} = 1.75$, $p = .191$).

These results suggest that in WT mice, the stimulus-specific habituation of DA signals to the LED observed after a 30s inter-stimulus interval (Figure 4) is no longer evident after 310s, consistent with a habituation response which then dishabituates [11]. In contrast, the stimulus-specific habituation to the LED in *Gria1*$^{-/-}$ mice that was not present after a 30s inter-stimulus interval (Figure 4) appears now to show stimulus-specific sensitization after a 310s inter-stimulus interval.

Overall, this pattern of differences was supported by a significant Genotype x HouseLED x SameDifferent interaction ($F_{1,\ 20.39}$ = 10.92, p = 0.003), that reflected a significant HouseLED x SameDifferent interaction in *Gria1*$^{-/-}$ mice ($F_{1,12.91}$ = 6.11, *p* = 0.028) but not in WT mice ($F_{1,\ 12.42}$ = 1.96, *p* = 0.186). There was also a significant main effect of Time ($F_{1,\ 21.45}$ = 87.72, p < 0.001), a HouseLED x Time ($F_{1,\ 57.08}$ = 7.53, p = 0.008) and a HouseLED x SameDifferent ($F_{1,\ 20.39}$ = 4.35, *p* = 0.05) interactions (all remaining effects Fs < 2.81, *p* > 0.099).

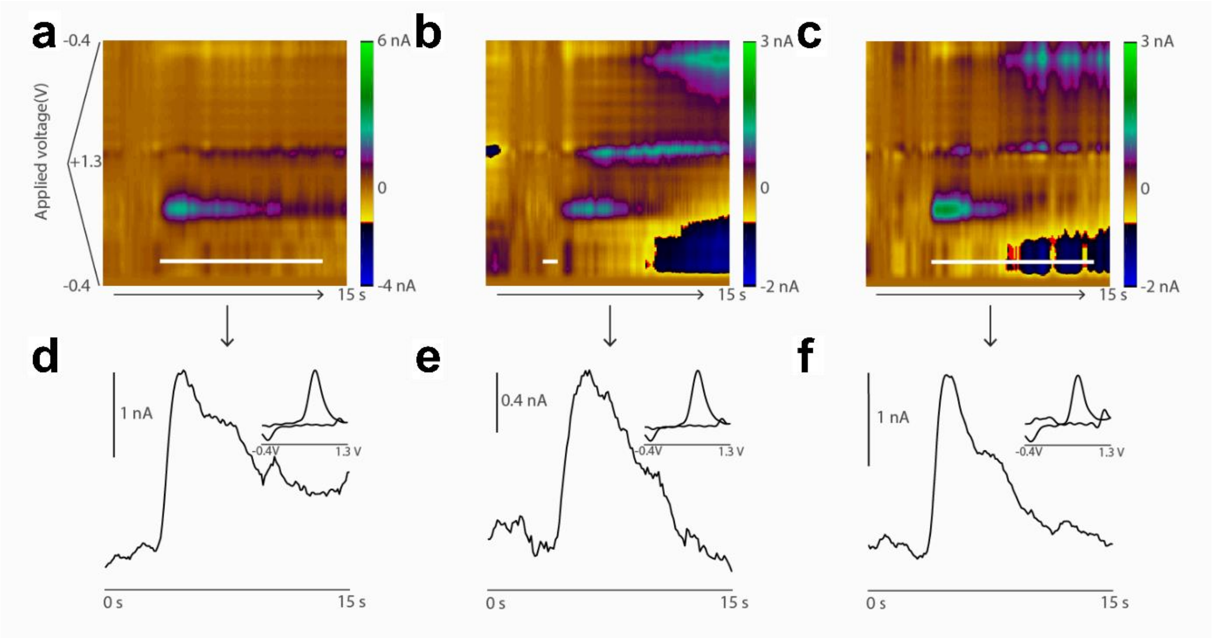


**Supplementary Figure 5. Representative dopamine responses to rewards and neutral light stimuli**

Representative single trial DA response in NAcc in response to a 10s LED light stimulus **(A,D)**, and a 22 µL sucrose reward **(B,E)** in the same WT mouse (from Fig2A and Fig 3B)., **(C,F)** DA release in response to the first presentation of a neutral 10s light stimulus, in a mouse completely naïve to reward delivery and the light in the test chamber i.e. DA response to a neutral stimulus when there is no reward expectation. Colour plots **(A-C)** show the background subtracted voltammograms as a function of applied voltage over time. White bar represents the duration of the light or reward stimulus. **(D-F)** Changes in DA levels (nA) over time, corresponding to the colour plots above. Inset, example cyclic voltammograms identifying the detected dopamine.

### Supplementary Discussion

#### GluA1-dependent plasticity mechanisms in the hippocampus underlie short-term memory

The role of GluA1 in hippocampal long-term potentiation (LTP) is well established. This mainly derives from hippocampal slice experiments in GluA1-KO mice (the same mice as used in the present study). Hippocampal LTP is significantly reduced or blocked altogether in GluA1-KO mice, although depending on the LTP induction protocol that is used there is sometimes a residual GluA1-independent form of LTP [12–16]. The role of GluA1 in hippocampal LTP is widely thought to reflect the rapid trafficking of GluA1-containing AMPA receptors into the post-synaptic density following NMDAR activation in order to increase the efficacy of synaptic connections [17, 18]. Consistent with this possibility Whitlock et al., [19] have shown an increase in GluA1 levels in the post-synaptic density of hippocampal neurons following a learning experience, an increase which lasted approximately 2 hr, and thus reflected a short-term memory trace in the hippocampus. These results are consistent with a role for GluA1 in experience-dependent synaptic plasticity in the hippocampus.

Furthermore, direct evidence for the role of GluA1 in experience-dependent synaptic plasticity comes from *in vivo* electrophysiological studies in GluA1-KO, in which neuronal activity can be measured directly in behaving animals. Notably, in a key study, Resnik et al., [20] conducted hippocampal single unit recording studies in wild-type and GluA1 knockout mice as they traversed a simple, linear maze track. Spatial rate maps emerged clearly and systematically with experience on the maze in WT mice on a trial-by-trial basis, whereas these representations failed to develop systematically within a session and were unstable in GluA1 KO mice. Thus, GluA1 knockout prevented experiential plasticity and the development of relevant cell assemblies in the hippocampus following experience on a simple maze task.

Finally, in a number of previous studies, we have also demonstrated the causal necessity of hippocampal GluA1 for short-term memory performance. Reinserting hippocampal GluA1 into GluA1-KO mice [e.g. 21] recovers short-term memory deficits in GluA1-KO mice. Conversely, ablation of hippocampal GluA1 in WT mice [22] is sufficient to cause the same short-term memory deficits observed in GluA1-KO mice. Taken together, hippocampal GluA1-dependent plasticity is an important cellular mechanism underlying short-term memory performance.

12. Erickson MA, Maramara LA, Lisman J. A single brief burst induces GluR1-dependent associative short-term potentiation: A potential mechanism for short-term memory. J Cogn

Neurosci. 2010;22:2530–2540.
